# A monkeypox mRNA-lipid nanoparticle vaccine targeting virus binding, entry, and transmission drives protection against lethal orthopoxviral challenge

**DOI:** 10.1101/2022.12.17.520886

**Authors:** Alec W. Freyn, Caroline Atyeo, Patricia L. Earl, Jeffrey L. Americo, Gwo-Yu Chuang, Harini Natarajan, Tiffany Frey, Jason Gall, Juan I Moliva, Ruth Hunegnaw, Guha Asthagiri Arunkumar, Clinton Ogega, Arshan Nasir, Hamilton Bennett, Joshua Johnson, Michael A. Durney, Guillaume Stewart-Jones, Jay W Hooper, Tonya Colpitts, Galit Alter, Nancy J. Sullivan, Andrea Carfi, Bernard Moss

## Abstract

Monkeypox virus (MPXV) caused a global outbreak in 2022, fueled by behaviorally-altered and enhanced human-to-human transmission. While smallpox vaccines were rapidly deployed to curb spread and disease among those at highest risk, breakthrough disease was noted after complete immunization. Given the imminent threat of additional zoonotic events as well as the virus’ evolving ability to drive human-to-human transmission, there is an urgent need for the development of a MPXV-specific vaccine that is able to also confer broad protection against evolving strains and related orthopoxviruses. Here, we demonstrate that an mRNA-lipid nanoparticle vaccine encoding a set of four highly conserved MPXV surface proteins involved in virus attachment, entry and transmission can induce MPXV-specific immunity and heterologous protection against a lethal vaccinia virus (VACV) challenge. Compared to Modified Vaccinia Virus Ankara (MVA), which forms the basis for the current MPXV vaccine, mRNA-vaccination generated superior neutralizing and cellular spread-inhibitory activities against MPXV and VACV as well as greater Fc-effector Th1-biased humoral immunity to the four MPXV antigens and the four VACV homologs. Single MPXV antigen mRNA vaccines provided partial protection against VACV challenge, while combinations of two, three or four MPXV antigen expressing mRNAs protected against disease-related weight loss and death. Remarkably, the cross-protection by multivalent MPXV mRNAs was superior to the homologous protection by MVA, associated with a combination of neutralizing and non-neutralizing antibody functions. These data reveal robust protection against VACV using an mRNA-based vaccine targeting four highly conserved viral surface antigens, linked to the induction of highly functional antibodies able to rapidly control viral infection.

## Introduction

First identified in the late 1950s in non-human primates(*1*), monkeypox virus (MPXV) caused intermittent, largely self-limiting, outbreaks in humans across central and west Africa following the cessation of smallpox vaccination(*2*). In 2003, monkeypox broke out in the US following the importation of wild rodents, infecting over 70 individuals, and resulting in the hospitalization of a young child(*3*). However, as with previous outbreaks, inefficient human-to-human spread led to rapid containment of the outbreak(*4, 5*). The current 2022 MPXV global outbreak led to infections in over 29 countries, resulting in approximately 80,000 infections and 52 deaths(*2, 6-8*), fueled by a combination of ease of travel and enhanced human-to-human transmission(*9*), calling for urgent public health action.

Smallpox vaccines, currently based on the live-attenuated Modified Vaccinia Ankara (MVA) strain, have been shown to attenuate MPXV disease both in epidemiologic studies and animal models(*10-15*). Thus, stockpiled JYNNEOS, an MVA-based smallpox vaccine, was rapidly deployed to induce immunity and attenuate disease in those at highest risk. Although data are still limited, breakthrough MPXV disease has been noted in the immediate period after both the first and second vaccine doses across vaccinated populations(*16*), pointing to incomplete protection afforded by the vaccine. Additionally, there were concerns about vaccine availability and challenges to manufacture additional JYNNEOS doses quickly. Moreover, given the extensive animal reservoir occupied by MPXV and its potential for continued evolution to cause enhanced human-to-human transmission, the development of a vaccine able to effectively limit MPXV, and potentially additionally orthopoxviruses, is urgently needed.

Immune correlate analyses have pointed to a critical role for the humoral immune response in protection against orthopoxviral infections. Specifically, mice deficient in B cells exhibit severe disease after infection, despite the presence of robust CD8+ T cell immunity(*12, 17, 18*). Similarly, in a non-human primate (NHP) model smallpox vaccine study, depletion of B cells, but not T cells, resulted in breakthrough infection after MPXV challenge(*12*). Vaccinia Immune Globulin (VIG), generated by pooling vaccinee plasma, was shown to prevent infection in close contacts of individuals with smallpox as well as help treat individuals with vaccine-related complications(*19*). However, in a study in cancer patients treated with hyperimmune antibody showed that protection against smallpox was strictly dependent on the activation of complement, suggesting that antibody binding alone may be insufficient for complete protection(*20*). Moreover, *in vitro* neutralization does not always correlate with protection *in vivo(21)* and vice versa(*22*) depending on the VACV antigen. Linked to the emerging appreciation for the critical role for Natural Killer (NK) cells in control of MPXV infection in mice(*21*), and the large number of genetic elements found in the poxviral genome used to evade complement(*22*), these data collectively argue for a key role for both neutralizing and non-neutralizing antibody functions in the control and clearance of the virus and in attenuation of disease.

Next generation vaccine discovery, aimed at generating highly efficacious, less reactogenic vaccines, have focused on the identification of specific sets of orthopoxviral immunogens for the design of recombinant vaccines(*15, 23-25*). Immunogenicity profiling of vaccine-induced immunity to MPXV has pointed to a set of surface proteins that are highly conserved across orthopoxviruses, and when targeted can block viral infection *in vitro* and confer immunity *in vivo(15, 24-26)* in a Th1 dependent manner(*27*). This immunogen target set includes proteins on the surface of the two infectious forms of orthopoxviruses: the intracellular mature virion (MV) and the extracellular enveloped virion (EV). Specifically, VACV homologs of MPXV antigens M1 and A29, involved in cellular entry on the MV, and A35 and B6 involved in transmission on the surface of the EV, provided complete protection from disease following MPXV challenge in both mice and NHPs(*12, 15, 23, 24, 28*). Given our emerging appreciation for the highly functional Th1-biased humoral immune responses induced by mRNA immunization(*29, 30*), here we aimed to test whether mRNA-based vaccination with these four highly conserved MPXV proteins could confer equivalent protection against orthopoxvirus infection as compared to MVA, which induces an immune response to a much larger array of proteins(*31-33*). We report that mRNA-lipid nanoparticle vaccination induced superior neutralizing activity compared to MVA and strong functional humoral immune responses across all four MPXV antigens and their VACV homologs. Moreover, the 4-antigen expressing mRNA conferred complete protection against death and morbidity following lethal VACV challenge while MVA-immunized animals experienced transient weight loss following challenge. Neutralizing and functional antibody responses were observed with both low and high mRNA dosing, linked to near complete protection against morbidity. Moreover, individual M1 or B6 expressing mRNAs conferred substantial protection against VACV, whereas vaccines containing a combination of mRNAs provided near sterilizing immunity against lethal challenge. Both neutralizing and Fc-profiles were strongly associated with reduced weight loss pointing to a critical role for both neutralizing and non-neutralizing cross-reactive antibody mediated protection against orthologous VACV challenge. Thus, MPXV-mRNA vaccines targeting even a single antigen can confer broad orthopoxvirus immunity, with combinations of MPXV providing near sterilizing immunity, comparable, if not superior, to homologous MVA immunization.

## Results

### Designed of modified membrane-bound MPXV antigens exhibit high surface expression

Previous studies have demonstrated that *in vivo* protection of animals can be achieved with immunization comprising just 2 to 4 conserved vaccinia virus (VACV) proteins, that are required for infection(*12, 25, 31, 32, 34, 35*). Given the complex life-cycle of orthopoxviruses, the inclusion of 1 to 2 antigens from the MV (A27 and L1) and 1 to 2 proteins from the EV (B5 and A33) provides protection against both forms of the virus, that have completely different surface antigens, and are both capable of causing infection. Thus, the four monkeypox (MPXV) orthologues (A29, M1, B6 and A35), which share 94.6%, 98.4%, 96.5%, and 95% amino acid identity to the VACV antigens (**Supplemental Figure 1**), were selected for vaccine design. Importantly, while the EV proteins B6 and A35 have signal peptides, enabling traffic through the ER during virus infection, the MV protein M1 traffics through the cytoplasm to insert into the viral membrane, and A29 binds to a viral transmembrane protein. Thus, similar to previous studies that observed increased immunogenicity with a cell-surface expressed VACV homolog of M1, here we developed plasma membrane-bound monkeypox immunogens (B6, A35, M1) with high surface expression in mammalian cell. Specifically, sequences for each of the 4 antigens were engineered to include a signal peptide, N-linked glycosylation sites were re-engineered, and/or transmembrane/cytoplasmic tails were modified. For B6, we tested the full-length mRNA sequence as well as an engineered version with a cytoplasmic tail (amino acids after residue 303) truncation. For A35, we tested the full-length sequence as well as another version with the cytoplasmic and transmembrane region (first 59 amino acids) replaced with an N-terminal transmembrane domain from an influenza N2. For M1, we added a signal peptide from an influenza H1, removed all the *N*-linked glycosylation codons by mutating threonine/serine residue codons to alanines, and tested a cytoplasmic truncated antigen (amino acids after residue 208). Flow cytometric analysis of antigen-expressing cells using polyclonal sera raised against the respective proteins of VACV showed that the addition of the transmembrane/cytoplasmic region led to superior cell surface expression (**Supplemental Figures 2**). Thus, these membrain bound designs, together with a secreted version of A29 (with addition of a signal peptide from an influenza H1 and removal of all the *N*-linked glycosylation sequons and cysteines), were incorporated into our vaccine construct.

**Figure 1.**
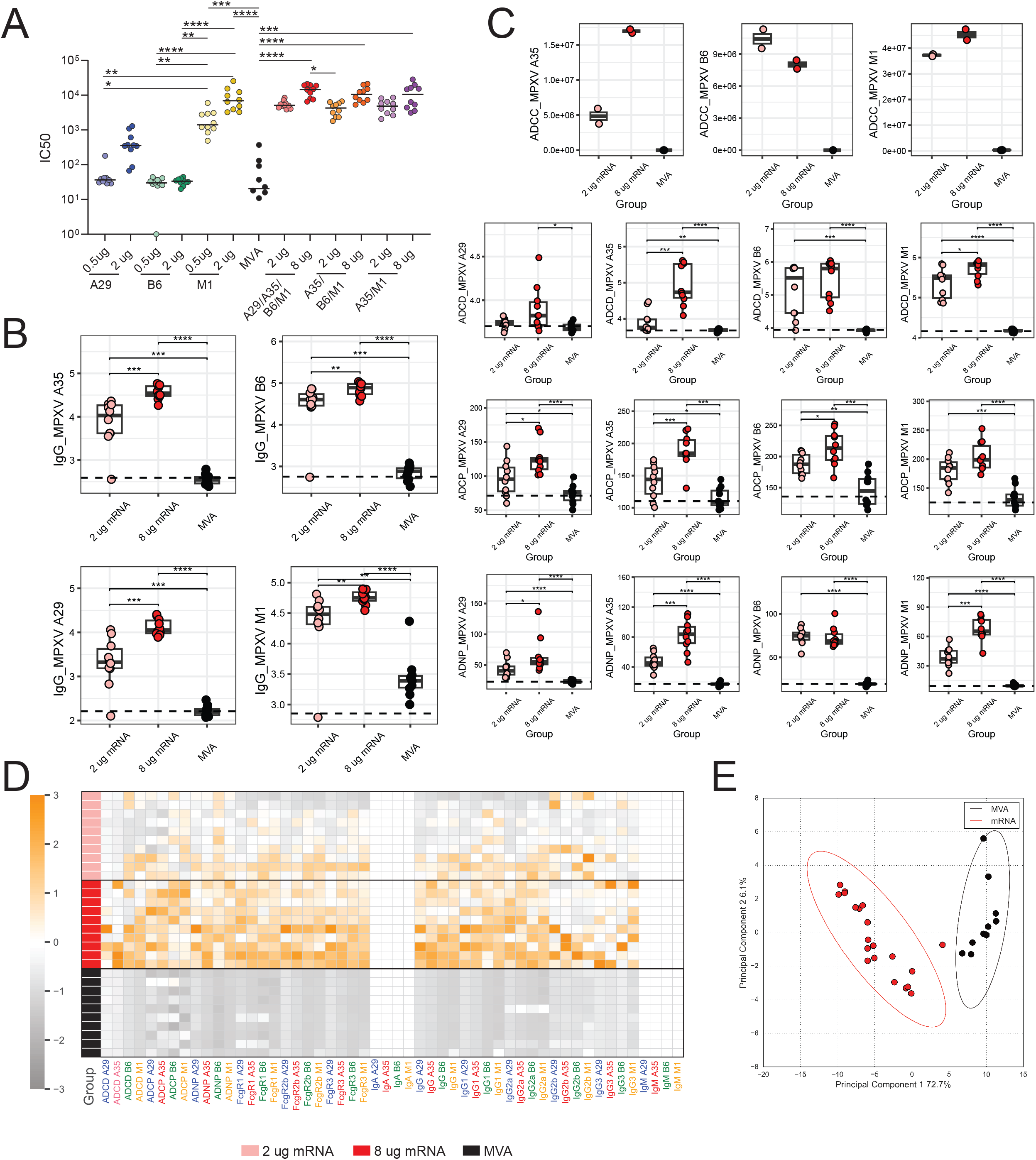
Distinct antibody signatures to A29, A35, B6, and M1 across mRNA and VACV immunized animals. a. The dotplots show the neutralizing antibody responses against MPXV-Z-1979 post-boost dose from a total of 160 mice divided into groups of 10. Significance was determined by a Kruskal-Wallis test, and comparisons were only made within single mRNA-immunized animals and against MVA or within combination mRNA-immunized animals and MVA. * p < 0.05, ** p <0.01, *** p<0.001, ****p<0.0001 b. The dot plots show the total IgG response, as measured by a multiplex Luminex assay, against the antigen listed. The dotted line represents the median response in the PBS vaccinated group. Significance was determined by a Mann-Whitney U test and corrected for multiple hypothesis testing using Benjamini-Hochberg method. * p < 0.05, ** p <0.01, *** p<0.001, ****p<0.0001. c. The dot plots show functional antibody responses: antibody dependent cellular phagocytosis (ADCP), antibody dependent neutrophil phagocytosis (ADNP), antibody dependent complement deposition (ADCD), and ADCC (ADCC). The dotted line represents the median response in the PBS vaccinated group. Significance was determined by a Mann-Whitney U test and corrected for multiple hypothesis testing using Benjamini-Hochberg method. * p < 0.05, ** p <0.01, *** p<0.001, ****p<0.0001. d. The heatmap shows the z-score antibody response for each animal. Orange indicates a higher response, whereas gray indicates a lower response. IgA and IgM were given a zero since all signal was at background. e. A principal component analysis (PCA) was built using the antibody profiles measured after mRNA vaccination and MVA vaccination. The red dots represent the individual antibody profiles in the mRNA group, whereas the black dots represent the individual vaccine response in the MVA group. The ellipses represent the 95% confidence interval for each group.

**Figure 2.**
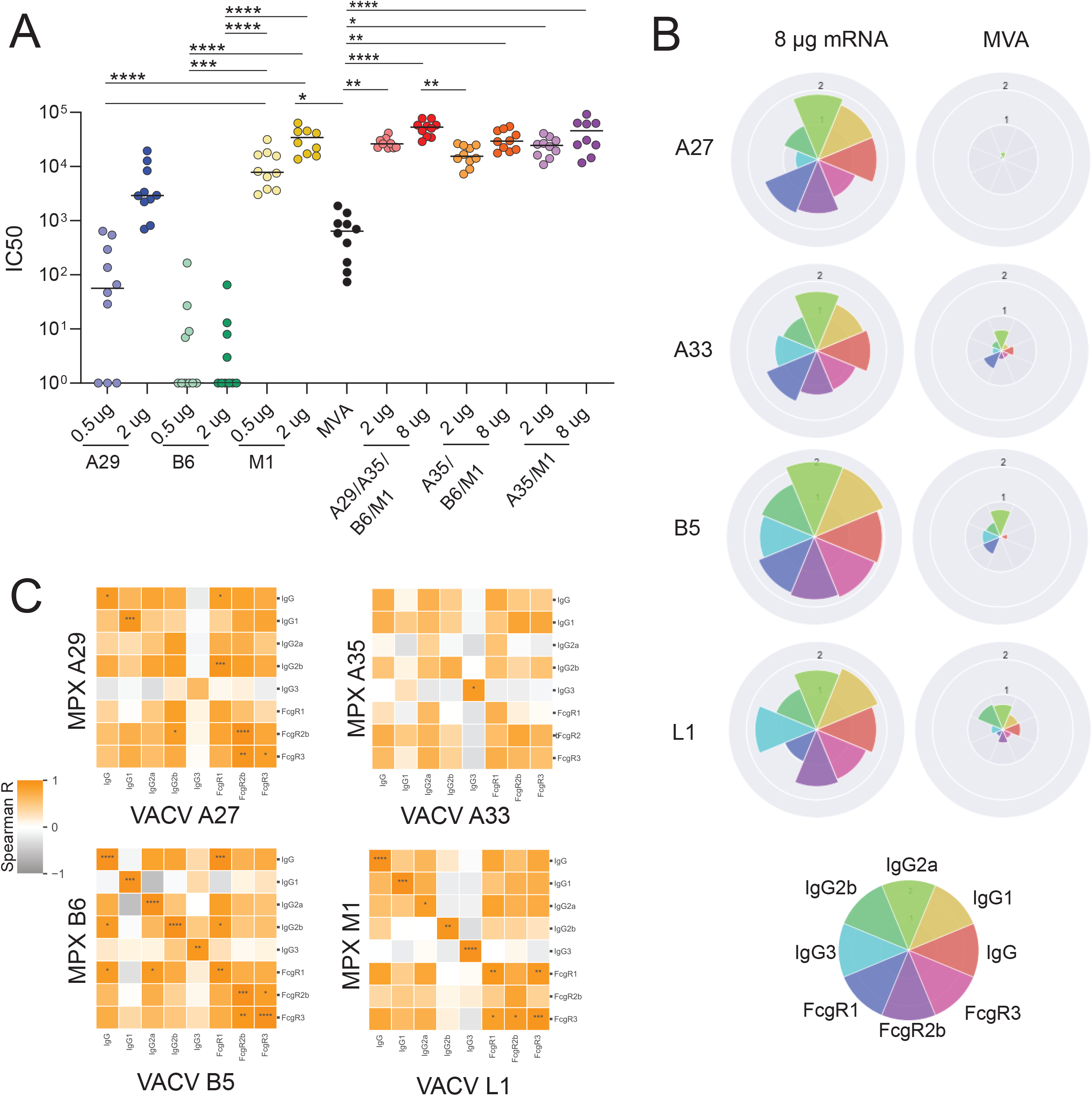
Substantial cross reactivity across MPXV and VACV antigens associate with mRNA dose and MVA induced protection against lethal VACV challenge. a. The dotplots show the neutralizing antibody responses against VACV-WR post-boost dose from the same 160 mice of Figure 1. Significance was determined by a Kruskal-Wallis test, and comparisons were only made between within single mRNA-immunized animals and against MVA or within combination mRNA-immunized animals and MVA. * p < 0.05, ** p <0.01, *** p<0.001, ****p<0.0001 b. The polar plots show the mean percentile rank for reach VACV feature for the 8 ug mRNA vaccine group and the MVA group. The color of the pie-slice represents a distinct IgG subclass or Fcg-recepror binding level shown in the legend, bottom right. The plots showing the response for the same antigen are comparable. c. The heatmaps show the spearman correlation of responses between MPXV and VACV orthologues in the 8 ug mRNA vaccine group. * p < 0.05, ** p <0.01, *** p<0.001, ****p<0.0001.

### MPXV mRNAs induce higher neutralizing titers and antibody profiles compared to MVA

Nanoparticles containing individual or combinations of A29, M1, B6 and A35 mRNAs were injected intramuscularly (IM) into BALB/c mice and again 3 weeks later, in a 2 dose regimen. As a comparator, a set of mice were immunized twice with 10 plaque forming units (PFU) of MVA, the attenuated smallpox vaccine that forms the basis of the approved JYNNEOS vaccine. The mice were bled at weeks 3 and 5 and the sera were analyzed for their ability to neutralize MPXV MVs that express green fluorescent protein (GFP) using a previously described quantitative high throughput flow cytometry assay(*36*). For animals that received 0.5 or 2.0 μg of mRNA encoding single MPXV antigens, the highest neutralizing antibody titer was observed with the M1-expressing mRNA, significantly surpassing neutralizing antibody responses observed with A29 and MVA (**Figure 1A**). Notably, appreciable neutralizing antibody responses were also detected after vaccination with the lower dose M1 that were superior in the neutralizing antibody levels induced by MVA, albeit at reduced levels compared to neutralizing levels in the higher dose group. Since the EV proteins B6 and A35 do not induce MV neutralizing antibodies, we next aimed to ensure that the inclusion of these antigen would not compete or reduce the neutralizing antibody levels that could be induced by M1 and A29. Thus, mice were vaccinated with combinations of mRNAs that included M1 and one or both of the EV proteins A35 and B6, totaling 2 μg for the low dose and 8 μg for the high dose. Again, neutralization of MPXV was dose dependent and B6 and A35 did not exhibit any interference, and the mice that received the mRNA vaccines induced higher neutralizing activity against MPXV compared to MVA (**Figure 1A**).

Using Luminex, we next confirmed the induction of binding antibodies across mice immunized with mRNA to the four MPXV target antigens (**Figure 1B**). Across the mRNA vaccine doses, larger differences were noted in the total IgG titers induced by the 2 μg and 8 μg mRNA groups for the A29 and A35 antigens than for M1 and B6 (**Figure 1B**). The nearly undetectable binding of IgG against these 4 antigens from MVA-immunized mice suggested that MVA-induced neutralization is likely directed at a larger number of distinct antigenic determinants(*15, 31-33*) and not exclusively to A29, M1, A35 and B6.

Next, we examined the overall MPXV-antigen specific antibody isotype and Fcg-receptor (FcgR) binding profiles induced by the quadrivalent mRNA given at the high or low dose compared to MVA. Little to no IgM and IgA were noted at peak immunogenicity to the 4 vaccine antigens across all 3 vaccine groups (**Supplemental Figure 3**), likely due to robust class-switching and non-mucosal site of vaccination. Conversely, distinct and diverse IgG subclass selection profiles were observed across the vaccine groups against the 4 antigens. A strong Th1-dependent IgG2a response was noted, with lower IgG2b, IgG1 and IgG3 responses. Similar IgG2a levels were noted against M1 and B6 across the 2 μg and 8 μg dose groups, whereas the 2 μg immunized animals induced lower levels of IgG2a-antibodies against A29 and A35. This highly functional IgG subclass bias was accompanied by a parallel FcgR binding profile, marked by enhanced FcgR binding antibodies in mRNA immunized animals, across the high affinity FcgR1, the opsonophagocytic FcgR2b, and cytotoxic Fcg3R (**Supplemental Figure 3**). Similarly, this IgG subclass skewing was also accompanied by the induction of humoral immune responses able to induce robust Fc-dependent effector functions, including antibody dependent cellular phagocytosis (ADCP), antibody dependent neutrophils phagocytosis (ADNP), antibody dependent complement deposition (ADCD), and antibody dependent cellular cytotoxicity (ADCC) (**Figure 1C**). These data highlight the Th1-biased humoral response, linked to robust FcgR binding and functions, elicited by mRNA vaccination (**Figure 1D**). Moreover, these data show that the humoral profile induced by mRNA vaccination is distinct from that induced by MVA vaccination (**Figure 1E**).

**Figure 3.**
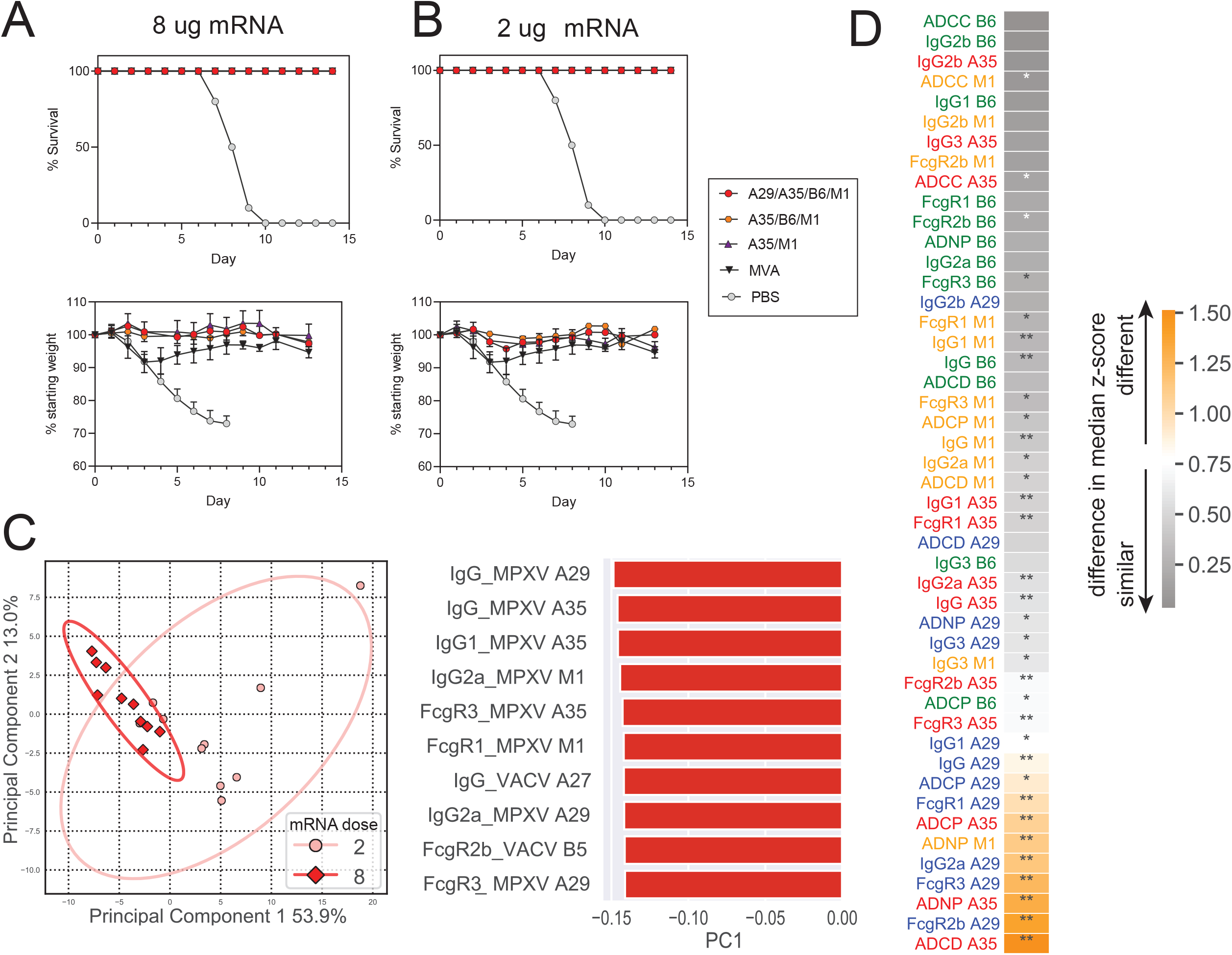
Dose independent conservation of B6 and M1 immunity. a. Survival and weight loss curves for the 8 μg bi-, tri- and quadri-valent mRNA vaccines, MVA, and PBS groups after lethal VACV challenge of the immunized mice described in Figure 1. Error bars represent SEM. b. Survival and weight loss curves for the 2ug bi-, tri- and quadri-valent mRNA vaccines, MVA, and PBS groups after lethal VACV challenge. c. The dot plot shows a principal component analysis (PCA) built on the antibody profiles in the 2 and 8 ug quadrivalent mRNA vaccine groups. Red diamonds show the antibody profile of individual animals in the 8 ug group, whereas pink circles show the antibody profile of individual animals in the 2 ug group. The ellipses show the 95% confidence interval for each group. The bar plot shows the features with the highest loading along principal component 1, all of which are enriched in the higher dose immunized animals (in dark red). d. The heatmap shows the difference in median z-score for each MPXV-specific antibody feature between the quadrivalent mRNA immunized groups. Gray indicates less difference in median z-score, whereas orange indicates higher difference in median z-score between the two groups.

### MPXV mRNAs induce cross-reactive antibodies to VACV antigens

Given the high degree of sequence conservation across MPXV and VACV (**Supplemental Figure 1**), we next sought to determine whether MPXV mRNAs induced cross-reactive antibodies to their VACV orthologues. Serum from mice that received individual and combinations of the 4 MPXV mRNAs were analyzed for their ability to neutralize VACV strain WR expressing GFP. The results were similar to that obtained with MPXV, marked by higher, but dose-dependent neutralizing titers at both 0.5 and 2 μg doses of mRNA vaccine containing the M1 antigen alone or in the 2 and 8 μg combinations of mRNAs, compared to MVA (**Figure 2A**). These neutralizing antibody responses were linked to the induction of a cross-reactive VACV-specific highly functional IgG2a-biased humoral immune response, across the 4 antigens, at higher levels than observed with MVA immunization (**Figure 2B**). Additionally, mRNA vaccination also induced strong and cross-reactive FcγR-binding responses across the VACV orthologoues (**Figure 2B**) across both dose groups (**Supplemental Figure 4**). Total IgG, IgG2a and FcγR-binding VACV and MPXV-specific responses were highly correlated, across the mRNA vaccine arms (**Figure 2C**). IgG3 and IgG2b, that were induced at lower levels (**Supplemental Figure 4**) were less well correlated pointing to mRNA induced antibody breadth across orthopoxviral protein orthologues (**Figure 2C**). Thus these data suggest that mRNA vaccination induced robust functional IgG cross-reactive neutralizing immunity that could potentially confer protection comparable to MVA against VACV.

**Figure 4.**
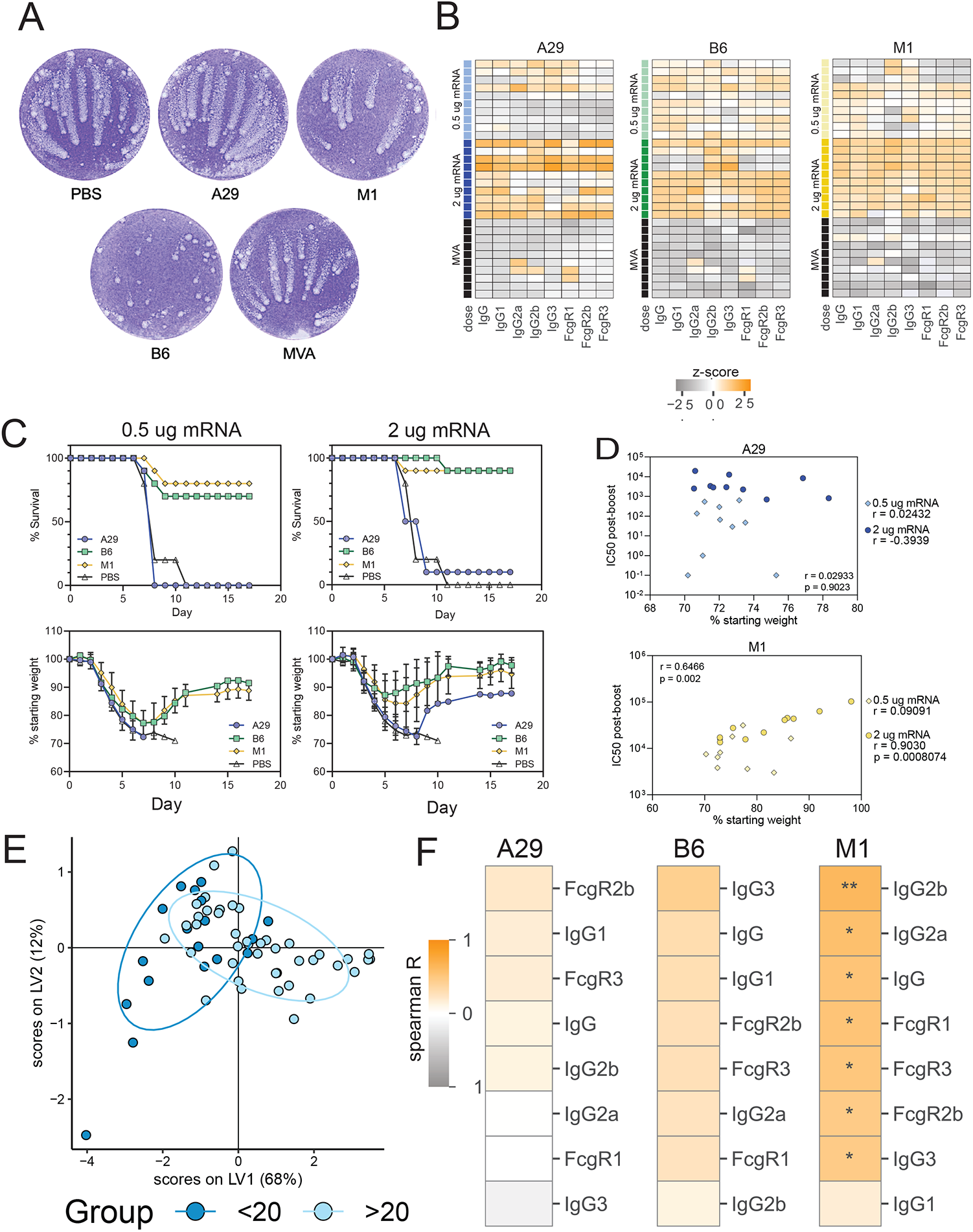
mRNA vaccine induced protection persists with combinations of as few as 2 antigens. a. The images of crystal violet stained B-S-C-1cell monolayers show the effects on VACV spread of addition of equal volumes of pooled sera from mice primed and boosted with the indictated individual mRNAs or MVA compared to control PBS. Serum was added 2 h after infection and incubation was continued for 40 h. Comet-like satellite plaques are due to spread of EVs. b. The heatmaps show the univariate levels of A29, B6, M1-specific IgG subclass binding induced for the vaccine antigens in the 2 μg an 8 μg single antigen mRNA immunized animal groups c. The plot shows the survival curves and percent of starting weights after VACV challenge of animals that received 0.5 ug mRNA dose (left) or 2 ug mRNA dose (right) of an individual MPXV mRNA. Error bars represent SEM. d. The plot shows the correlation of neutralizing antibody to percent starting weights of mice immunized with 0.5 or 2.0 μg of mRNA encoding A29 or M1. Correlation coefficients (r) and significance are shown. e. The PLSDA shows the multivariate discrimination between animals that lost more than 20% body weight (light blue) and less than 20% body weight (dark blue). f. The heatmap shows the spearman correlation of each feature against A29, B6, or M1 versus the minimum percent weight of the matched-antigen vaccinated animals from starting weight after challenge. Orange inidicates less weight loss, whereas gray indicates a greater change in weight.

### Low and high doses of MPXV mRNA combinations protect mice challenged with a lethal dose of VACV

Given the homologous nature, broader range of antigens, and native antigen expression by MVA, it may be expected that MVA would have an advantage over MPXV mRNAs in a vaccinia virus challenge model. To test this hypothesis, the BALB/c mice that had been immunized with MVA or or two different doses of bi-, tri-, or quadrivalent mRNAs were challenged with a lethal 10^6^ PFU dose of VACV WR via the intranasal route 3 weeks after boost vaccination. Severe weight loss was observed in non-immunized animals and all succumbed to disease within 7 to 10 days (**Figure 3A-B**). As expected, all MVA immunized mice survived lethal challenge, despite a transient early loss in weight (**Figure 3A-B**). In contrast, 8 μg mRNA immunized animals showed complete survival and no weight loss after challenge. Similarly, mice immunized with 2 μg of the MPXV-mRNA vaccine showed complete survival and only a slight transient weight loss after challenge (**Figure 3A and 3B**).

Given the robust neutralizing titers induced by bivalent (A35/M1) and trivalent (A35/B6/M1) vaccines against VACV (Figure 2A), we also determined whether MPXV bivalent (A35/M1) and trivalent (A35/B6/M1) mRNA vaccination could also afford protection against lethal VACV challenge. Remarkably, both vaccines conferred conplete survival (**Figure 3A and 3B, upper**), and nearly identical protection against weight loss as compared to the quadrivalent vaccine (**Figure 3A and 3B, lower**). These data point to robust, nearly sterilizing protection, afforded by the bi-, tri-, or quadri-valent MPXV mRNA vaccination against lethal VACV challenge.

### mRNA dose-dependent and -independent antibody responses

Despite the nearly dose-independent protection against lethal VACV challenge across the vaccines (**Figure 3A and B**), neutralizing and binding antibody profiles differed significantly across the vaccine doses (**Figure 1**). Thus, to begin to define whether particular antibody parameters were more highly conserved across doses, and thus potentially contributed to the similar levels of protection observed across vaccine doses, we next defined the antibody features that were most affected by vaccine dose. Focusing on the quadrivalent vaccine, a principal component analysis (PCA) clearly illustrated signficaint differencs in binding antibody profiles between the high and low dose mRNA vaccine groups (**Figure 3C, left**), driven largely by vaccine dose-dependent difference in A29L and A35R-specific binding antibody responses that were largely enriched in the 8 ug dose group (**Figure 3C, right**). However, to define the specific antibody features that were conserved across the mRNA dose groups, we plotted the difference in median z-score between the dose groups for all MPXV antibody features (**Figure 3D**). Specifically, we observed robust conservation of M1 and B6 antibody responses (minimal z-score differences, top, in grey), whereas significant differences were noted for A29 and A35 (bottom, in yellow), pointing to a potential dominant dose-independent role for M1 and B6 in cross-VACV protection.

### Distinct antigen-specific antibody protective correlates of immunity

M1 immunization, even alone, led to robust neutralizing antibody responses to MPXV and VACV mature virions and A29 immunization alone induced intermediate levels of neutralization (**Figure 1A and Figure 2A**). In contrast B6 immunization alone led to negligible levels of neutralizing antibodies, but importantly led to reduced spread of enveloped virus (**Figure 4A**). In contrast, sera from mice immunized with A29 had no effect on enveloped virus spread, and M1 and MVA sera had only a small effect compared to the PBS control (**Figure 4A**), pointing to an alternate mechanism of viral blockade induced by B6 that was more potent for the mRNA vaccine than MVA. Thus, despite the induction of high levels of binding IgG1 to the individual antigens in the higher dose group, significant differences in titers were noted across the antigens in the lower dose group, as well as across antibody subclasses and FcγR binding levels (**Figure 4B**), in addition to striking differences in the ability of these antibodies to prevent viral spread (**Figure 1A** and **Figure 4A**). Thus, to begin to define the protective cross-reactive mechanism of action of individual antigen-specific responses against lethal VACV challenge, the mice immunized with 0.5 or 2 μg of individual M1, B6, and A29 MPXV mRNAs (described in **Figures 1 and 2**) were challenged. All animals lost weight but the majority of those that received B6 or M1 mRNA recovered (**Figure 4C, lower**). A dose effect for B6 and M1 was observed as significantly less weight loss occurred on all days between 7 and 17 (median p=0.008 for B6 and p=0.02 for M1) of mice that received 2 μg of mRNA compared to 0.5 μg. In contrast to the protection seen with B6 and M1, all mice immunized with the low dose of mRNA to A29 succumbed to infection as did all but one receiving the high dose of A29 mRNA (**Figure 4C, upper**). Moreover, except for one outlier, the weight loss curve of mice receiving the high dose of A29 mRNA was superimposed on the weight loss curve of unimmunized mice (**Figure 4C, lower**). Thus, although both M1 and A29 mRNAs induced antibodies that neutralized MVs *in vitro* (**Figure 4A**), only M1 was protective when administered alone (**Figure 4B**). This result was not entirely unexpected as a similar disparity between *in vitro* neutralization and protection was previously reported using soluble VACV A27 for vaccination(*14*), although in DNA vaccine experiments A27 increased protection when combined with VACV A33 or B5(*32*). Furthermore, we observed no correlation between VACV neutralization (IC50) post-boost and the prevention of weight loss after challenge of mice immunized with the low or high doses of A29 (**Figure 4D, top**). Furthermore, A27 delivered as a DNA vaccine alone did not protect mice, but when combined with A33 or B5, the level of protection achieved by those targets alone was improved significantly. Moreover, while no correlation was observed in A29 neutrlizing activity and weight loss, a highly significant correlation was observed between neutralization and weight loss in mice immunized with the high dose of M1 (**Figure 4D**). Although B6 immunization did not induce MV neutralizing antibodies, the inhibition of EV spread (**Figure 4A**) can explain the protection nearly equivalent to M1. Thus, these data suggest that distinct antibody blocking mechanisms likely play a key role in protection against orthopoxviruses.

Despite the observation for distinct blocking functiosn of B6 and M1 antibodies, immunization with individual MPVX mRNAs did not afford sterilizing immunity. Rather post-infection control and resolution of disease was observed across both B6 and M1 immunized animals, suggesting that blocking alone is unlikely to be sufficient to explain post-infection control and clearance of the virus. To therefore define whether particular antibody features were highly associated with reduced weight loss, we performed a partial least squares discriminant analysis (PLS-DA) across all antigens. In this analysis, all singly-immunized animals were combined, and all antibody responses (neutralization, isotype, subclass, and FcR binding) were included into the model to define correlates of immunity against severity of disease. To perform the PLS-DA, the animals were split into two groups: a group of animals that lost greater than 20 percent of their starting body weight after challenge and a second group of animals that lost less than 20 percent of their starting body weight after challenge. The PLS-DA model showed that the two groups could be differentiated in multivariate space (**Figure 4E**, CV = 78%), suggesting that specific antibody profiles strongly predicted the degree of protection afforded by MPXV mRNA vaccination against VACV challenge induced weight loss. To more accurately define the antibody features associated with protection against disease, we performed a regression analysis by calculating the Spearman correlation of each antibody feature against maximum percent body weight after challenge (**Figure 4F**). This analysis showed that neutralizing antibody levels and class switched highly functional IgG2b antibody levels were among the strongest correlates of immunity against weight loss in M1 immunized animals (**Figure 4F**). Given the high affinity of IgG2b for FcgR4, involved in macrophage and neutrophil opsinophagocytic activity(*37*), as well as its more constrained T like structure that has been proposed to be important in the recognition and control of mucosal pathogens(*38*), these data point to a critical role for both the antibody antigen binding domain (Fab) and constant domain (Fc) qualities in protection against VACV induced disease. Moreover, additional IgG subclasses and FcgR binding levels were significantly associated with reduced weight loss in M1 immunized animals, after correction for multiple comparisons, supporting a role for Fc-functional antibody mechanisms in protection against disease. IgG subclass and FcgR binding were associated with reduced weight loss in B6 immunization animals, although these relationships did not survive multiple comparisons, potentially due to limited animal numbers. Simiarly, after multiple comparisons, a correlate of immunity was not observed in A29 mRNA immunized animals, potentially related to the limited protection afforded by this antigen alone. However weak correlations were observed only for FcgR-binding and IgG subclass levels, pointing to a potential collaborative role for both cross-reactive binding and Fc-recruitment in the control and clearance of the VACV following challenge. Collectively, these data suggest that neutralizing and non-neutralizing antibody functions likely provide orthogonal mechanisms of immune control, potentially offering first and second-line immune protection against MPXV and future related orthopoxviruses.

## Discussion

Since the development of the smallpox vaccine(*39*), using cowpox-like virus as the immunogen, three generations of the related VACV-based-smallpox vaccines have been developed(*1, 40, 41*). However, over the past decade, a large number of newer vaccine platforms have emerged that are able to drive robust humoral and cellular immunity(*42, 43*), including the mRNA vaccine platform, able to rapidly and flexibly deliver pathogen-antigens to the immune system to drive immunity. While attenuated pathogens induce immunity against the complete pathogen, current mRNA vaccination only delivers components of the target pathogen. Previous studies focused on key poxviral targets required for viral infection had shown similar protection compared to whole attenuated vaccination in both mice and NHP(*23-25, 32, 44*). Thus, here we tested whether mRNA delivery of a set of just 4 highly conserved antigens, involved in viral binding, entry and transmission expressed on the outer membrane of the mature virion (MV) or the enveloped virion (EV), could confer protection comparable to MVA.

Identification of key antigens, critical for vaccine development, is complicated by both the large size of the orthopoxviral genome(*45-49*), as well as the diversity of genes, many of which remain poorly functionally defined across these viruses. Moreover, significant variation exists among the orthopoxviruses, with much of the variation largely in virulence factors encoded on the genomic end regions of the viral genome, although the central region of the genomes exhibit high sequence identity(*50-52*). The proteins known to be involved in infection, both in the mature and enveloped virus, are remarkably conserved across VACV and MPXV, and the VACV proteins have been shown to be immunogenic and confer protection against VACV(*24, 25, 27, 28, 41, 53*). Because the mature and the enveloped viruses have distinct roles in infection, the inclusion of antigens from both viral forms is necessary for complete protection in animal models(*23, 24*). Moreover, vaccination with these proteins has also been shown to confer protection against additional orthopoxviruses such as ectromelia virus and rabbitpox(*54, 55*) and rabbitpox virus(*56*). Likewise, here immunization with MPXV derived M1, A29, A35 and B6 led to cross-reactive IgG responses to VACV A27, A33, B5, and L1. Similarly, human monoclonal antibodies to A27, A33, B5, and L1 have been shown to limit MPXV infection, and have been discussed as potential cross-reactive therapeutics for prophylactic use(*57*). Moreover, the cross-reactive responses exhibited comparable neutralization and FcgR binding profiles. Surprisingly, while both MVA and our 4-valent MPXV vaccine both conferred complete protection against lethal VACV challenge, superior protection against weight loss was observed using the orthologous MPXV mRNA immunogen, even at a quarter of the higher dose. Furthermore, even a tri- and bi-valent vaccine conferred near sterilizing immunity against orthologous VACV challenge, and single-valent (M1 or B6) mRNA vaccines also conferred protection against death, although M1 induced potent neutralizing antibody to the MV form of VACV and B6 prevented spread of the EV form. Thus, it is likely that the highly functional, cross-reactive responses induced via vaccination, to conserved antigens on the MV and EV particles, are likely to confer multi-functional (neutralizing and Fc-effector) immunity able to both block and rapidly eliminate viral infection across multiple orthopoxviruses.

In the absence of non-survivors following challenge with the quadrivalent vaccine, immune correlate analyses following vaccination in the lethal VACV challenge model have been complicated. Previous immune depletion studies, including the depletion of T and B cells have clearly illustrated the critical role of antibodies in protection against infection(*12*). Yet, the precise mechanisms by which antibodies confer protection are incompletely defined. Antibodies to the 4 target proteins selected for the mRNA vaccine either neutralize MV (M1 and A29), reduce cell-to-cell spread of EVs (A35 and B6) *in vitro*, or may drive effector-cell mediated clearance of infected cells or virus *in vivo*. Furthermore, a mAb to the VACV homolog of M1 prevents virus entry into cells at the post-hemifusion step and antibodies to the VACV homolog of B6 aggregate EVs and prevent their binding to cells(*58-60*). The partial protection afforded by single antigen mRNA vaccines, in the setting of variable levels of *in vitro* neutralization, offered a unique opportunity to define correlates of immunity against orthopoxviruses. Specifically, a combination of neutralization and unique IgG subclasses with high affinity for FcgR binding were enriched among animals exhibiting reduced weight loss. Moreover, it is likely that all of the mRNA-vaccinated mice became infected upon challenge, as there was a slight transient drop in weight, though less than that of mice immunized with MVA. Although little or no IgA was produced in mice with systemic vaccination, IgG/FcgR reponses play a critical role in reducing infection and driving pathogen clearance in the lungs(*61, 62*). Given the abundance of innate immune cells in the lung(*63, 64*), complement deposition and NK cell functions were robustly maintained across the mRNA vaccine doses, and both have been previously implicated in protection in immunocompromised humans(*27, 45*), in mice(*23, 65*) or in the CAST mouse model(*21*). In an *in vitro* model enhanced protection by combinations of antibodies was shown to be due to complement mediated anti-A33 antibody-dependent disruption of the EV membrane exposing the inner MV membrane to neutralizing M1 antibody(*66, 67*). Thus, the data presented here point to a critical collaborative role for both neutralizing and non-neutralizing antibody functions in orthologous protection against VACV using an MPXV mRNA vaccine.

Here, we showed that an MPXV mRNA vaccine can induce robust neutralizing and functional cross-reactive antibodies able to confer comparable, if not superior, protection against a lethal challenge of an orthologous orthopoxvirus compared to MVA. Moreover, while as few as a single MPXV-antigen expressing mRNA was able to confer complete protection against lethal challenge, combinations of antigens conferred near sterilizing immunity against orthologous viral challenge. Linked to the unprecedented real-world data related to the impact of mRNA vaccination on limiting the COVID-19 pandemic(*43*), the application of this flexible platform in response to emerging and re-emerging pathogens, including MPXV, offers a unique opportunity to rapidly limit spread. Yet, while mRNA induced vaccination has been shown to drive broad and durable T cell immunity in the setting of COVID-19 vaccines(*68, 69*), whether MPXV mRNA-vaccination also induced T cell functions that may contribute to mRNA induced immunity is of great interest and could provide enhanced insights on the application of mRNA vaccines both prophylactically or therapeutically against MPXV, other orthopoxviruses, and beyond. However, whether responses induced by mRNA vaccination will maintain the lifelong durability of immunity that has been associated with current smallpox vaccination(*70, 71*) remains uncertain. Yet the flexibility and speed of mRNA vaccine design offers new opportunities for orthopoxvirus vaccine design, allowing for the rapid adaptation, production, and deployment of vaccines, in the absence of the adverse events that have been associated previously with smallpox vaccines.

## Material and Methods

### Vaccine Design

mRNAs encoding M1, A29, A35 and B6 were selected from clade II of MPXV, using sequences from MA001 (NCBI Accession No. ON563414), which was one of the first few completely sequenced genomes publicly available from the outbreak (**Supplemental Figure 1**). The mRNA Drug Substance was formulated in a mixture of four lipids [SM-102 (a novel, ionizable lipid); PEG2000-DMG; 1,2-distearoyl-snglycero-3-phosphocholine (DSPC); and cholesterol] at a total lipid content of 9.7 mg/ml, in 20 mM trometamol (Tris) buffer containing 87 mg/ml sucrose and 10.7 mM sodium acetate at a dosage strength of 0.2 mg/ml, at pH 7.5. The Drug Product was stored at -60°C to -90°C.

### Antigen characterization

Expression of all mRNA-encoded antigens was assessed using Expi293 suspension cells (Thermo) incubated at 37°C, 8% CO_2_, 125 rpm in an Infors Multitron incubator (**Supplemental Figure 2**). Cells were seeded at 1.10^6^ /ml and 5ml/well in 24 well plates for 48 h (membrane constructs) or 72 h (secreted constructs). Two doses of unformulated mRNA (500 ng/ml and 100 ng/ml) were transfected in parallel using a TransIT-mRNA transfection kit (Mirus). For analysis of membrane-bound antigen expression, cells were harvested, blocked, stained with antigen-specific primary rabbit polyclonal antibodies (NR-629 for B6R, NR-628 for A35R, NR-631 for M1, all sourced from BEI Resources) and an Alexa Fluor 647 labeled goat anti-rabbit IgG (Southern Biotech) secondary in addition to Aqua (BD Biosciences) to differentiate live and dead cells. Flow cytometry was performed on an iQue3 instrument (Sartorius) with a sip rate of 1.5 ul/s for 18s. Data analysis was performed using ForeCyt software (Sartorius) by gating successively on cells, singlets, live cells and positive cells to determine the positivity rates and mean fluorescence intensities for construct comparisons analyzed using Prism software (GraphPad). For analysis of secreted constructs for A29L, we utilized a Jess automated immunoassay system (ProteinSimple/Bio-Techne). Neat supernatants and biotinylated protein standards were pipetted into designated wells in instrument-specific sample plates for assay initiation. Samples were diluted with sample buffer, denatured with 400 mM DTT before capillary-based separation. Standards were detected with streptavidin-HRP (ProteinSimple/Bio-Techne) while A29L antigen was detected with rabbit polyclonal NR-627 primary (BEI Resources) and an anti-rabbit IgG-HRP (R&D Systems). Data analysis was performed using Compass software (ProteinSimple/Bio-Techne).

### Mouse Study

The study aimed at determining the ability of each vaccine composition to generate antibody responses, with further depth into functional responses, the observation of interference and dose effect on responses, and to determine the efficacy of the candidate vaccine in BALB/c VACV challenge model. Two dose levels (0.5 μg and 2 μg) of individual mRNAs and two dose levels (2 μg and 8 μg) of the bi-, tri-of quadrivalent vaccine were administered IM in a volume of 50 μl. The 10^7^ PFU of MVA was also administered IM in a volume of 50 μl. Immunizations were performed at week 0 and 3, in a total of 160 female BALB/c mice. Animals were then challenged with VACV WR strain via the intranasal route 3 or 5 weeks after boost vaccination at a challenge dose of one million plaque forming units. Serum was collected 1 day prior to the boost and three weeks after the boost for antibody assessment. Morbidity of mice after intranasal challenge with a lethal dose of VACV WR was captured as percent of starting weight determined on a daily basis. All were humanely euthanized after reaching 70% starting weight by day 8.

### Virus neutralization and spread inhibition assays

The rapid, sensitive and quantitative 96-well plate semi-automated, flow cytometric assay was carried out using VACV strain WR expressing GFP as previously described(*36*). Recombinant MPXV Z-1979 expressing GFP was constructed and used in a similar manner to assay MPXV neutralizing antibody. Several 2-fold dilutions of heat inactivated immune serum (56°C for 30 min) from individual mice were prepared in 96-well, round bottom polypropylene plates using spinner modified MEM containing 2% fetal bovine serum (Spinner-2%). Approximately 2.5×10^4^ PFU of WR- or MPXV-GFP expressing viruses was added to each well and plates incubated at 37°C for 1 h. After incubation, 10^5^ HeLa S3 cells were pipetted into each well and plates were incubated for an additional 16-18 h at 37°C. The cells were fixed in 2% paraformaldehyde and GFP expression measured and quantitated using a FACS Canto II flow cytometer and FlowJo software (BD Biosciences). IC50 values were calculated using Prism software (GraphPad/Dotmatics).

The spread comet inhibition assay was carried out using VACV strain IHD-J as described previously(*67*). BSC-1 cells grown in 12-well tissue culture plates were infected with 30 PFU VACV strain IHD-J for 1 h at 37°C after which input virus was removed and cell monolayers were washed 3x with EMEM supplemented with 2% fetal bovine serum (EMEM-2%). Upon aspirating the final wash, cells were overlaid with 0.75 ml EMEM-2% containing a 1:50 dilution of heat inactivated pooled immune serum from mice receiving a high dose prime and boost of A29 mRNA, M1 mRNA, B6 mRNA, 10^7^ PFU MVA, or control PBS. The cells were incubated with diluted serum for an additional 40 h at 37°C and then stained with crystal violet prior to visualization.

### Luminex

MPX-specific M1, A29, A35, B6, and VACV-specific L1, A27, A33 and B5 antigens were coupled via carboxyl chemistry to Magplex® fluorescently bar-coded beads (Luminex Corporation) with sulfo-NHS and EDC (Thermo Fisher) per manufacturer’s instructions. These beads were then incubated with diluted, heat inactivated serum samples for 2 hours at 37°C, shaking. To detect total IgG, IgG1, IgG2a, IgG2b, IgG3, and FcgR (FcgR1, FcgR2a, FcgR2b, FcgR3, FcgR4) binding, samples were incubated with beads at a dilution deterimed by previously run dilution curves. Each sample was assayed in duplicate. Beads were then washed to remove unbound sample and incubated with PE-labeled secondary detection reagents (anti-Ig isotypes from Southern Biotechnology and FcgRs from Sino Biological). Excess detection antibody was washed away, and samples were quantified on Flexmap 3D (Luminex Corporation).

### Antibody dependent cellular phagocytosis

MPX-specific M1, A29, A35, B6 were biotinylated with EZ-link Sulfo-NHS-LC-LC-Biotin (Thermo Fisher) and excess biotin was removed using a Zeba size exclusion column (Thermo Fisher). Biotinylated antigens were coupled to Neutravidin yellow-green fluorescent beads (Thermo Fisher). Antigen-coupled beads were incubated with 1:500 diluted serum samples for two hours at 37°C, then washed to remove unbound sample. J774-1 cells (ATCC) were added to immune complexes and incubated for 1 hour at 37°C. Cells were then washed to remove unbound antibodies, fixed, and quantified on the iQue 3 VBR using Forecyt software (Sartorius). Phagocytic scores were calculated by multiplying the percentage of bead positive cells by the bead fluorescence GMFI of bead positive cells and dividing by 100,000. Each sample was assayed in two independent replicates.

### Antibody-dependent neutrophil phagocytosis

A mouse ADNP assay was adapted from the previously published protocol(*72*). MPX-specific M1, A29, A35, B6-coated beads were created as for ADCP assay. Antigen-coupled beads were incubated with 1:500 diluted serum samples for 2 h at 37°C, then washed to remove unbound sample. Fresh primary mouse white blood cells were isolated from the bone marrow of C57BL/6 mice and incubated with immune complexes for 1 h at 37°C. Cells were then washed to remove unbound antibodies, labeled for CD66b (anti-human CD66b fluorescent antibody from Biolegend), fixed, and quantified on the iQue 3 VBR using Forecyt software (Intellicyt). Neutrophils were identified as CD11b+/Ly6G+ positive. Phagocytic scores were calculated by multiplying the percentage of bead positive cells by the bead fluorescence GMFI of bead positive cells and dividing by 100,000.

### Antibody-dependent complement deposition

ADCD was adapted from the previously published protocol(*73*). MPX-specific M1, A29, A35, B6-coated beads were created as for Luminex. Antigen-coupled beads were incubated with 1:1000 diluted heat inactivated serum samples for 2 h at room temperature, shaking. Plates were then washed to remove unbound antibody. Immune complexes were then incubated with reconstituted lyophilized guinea pig complement (Cedarlane) for 20 min at 37C, shaking, and excess complement was washed off. Immune complexes were stained with anti-guinea pig C3 fluorescent antibody (MP Biomedicals). Excess staining antibody was removed by washing and immune complexes were quantified on the iQue 3 VBR using Forecyt software (Intellicyt). Complement deposition was indicated by the C3 GMFI of immune complexes. Each sample was assayed in two independent replicates

### Antibody-dependent cell-mediated cytotoxicity reporter assay

Antibody dependent cellular cytotoxicity was quantified using the Promega Fc reporter system(*74*). Chinese hamster ovary-K1 cells (ATCC) were transfected with x μg of pCAGGS plasmid DNA expressing membrane-bound MPXV antigen M1, A29, A35, or B6. Cells were allowed to incubate at 37°C with 5% CO_2_ for 48 h. Serum was serially diluted in Roswell Park Memorial Institute media supplemented with 4% Ultra-Low IgG FBS (Assay buffer; Gibco). Assay buffer and diluted serum was added to transfected cells. Promega FcgRIV expressing Jurkat cells were diluted in warm assay buffer and added to transfected cells and serum for 6 h at 37°C with 5% CO_2_. BioGlo luciferase substrate was warmed to RT and added to each well. Plates were read immediately on a Pherastar FS plate reader (BMG Labtech). Data were analyzed using Prism 9 (GraphPad) and are reported as Area Under the Curve (AUC) after curve analysis using the [Inhibitor] vs. response -- Variable slope (four parameters) with Baseline set to three times the standard deviation of negative wells per plate.

### Analysis

Univariate analysis was performed in GraphPad Prism, version 9.0 and R studio, R version 4.2.2. Univariate plots show the average of two replicates for all assays. For neutralization, significance was determined using a Kruskal-Wallis test and significance is only shown for differences between the same timepoint. For antibody measurements, significance was determined by a Mann-Whitney U test and corrected for multiple hypothesis testing by the Benjamini-Hochberg method. A 5-point dilution curve was performed for ADCC, and an area under the curve was calculated in Prism. Principal component analysis (PCA), heatmaps, and polar plots were visualized in Python 3.11. PCA was performed using sklearn.decomposition.PCA package. Prior to PCA, missing Luminex values were imputed using k-nearest neighbors, Luminex and ADCD data was log10 transformed, and all data was centered and scaled. For heatmaps, the data was z-scored. All correlation heatmaps show spearman correlation. Significance for correlation heatmaps was determined using the scipy.stats.spearmanr function in python. For polar plots, the median log10-transformed MFI of the specific feature of the 8 ug mRNA or MVA group was corrected by subtracting the median log10-transformed MFI of the PBS group for the same feature. For concordance heatmaps, the data was z-scored, and the median of each group for each feature was determined. The heatmap shows the absolute value of the difference between the 8 μg and 2 μg mRNA groups for each feature. For significance on the heatmap, a Mann-Whitney U test was performed for each feature between the two groups to determine whether the feature was significantly different between the two groups. P-values were adjusted using the Benjamini-Hochberg method. Significance was plotted in the heatmap as a star: * p <0.05, ** p< 0.01, *** p<0.001, **** p <0.0001.

Partial-least squares discriminant analysis (PLSDA) was performed in R (version 4.2.2.). The systemseRology R package (v1.0) (https://github.com/LoosC/systemsseRology) was used for PLS-DA, with few changes. Briefly, the data was log10 transformed, centered and scaled. Least absolute shrinkage and selection operator (LASSO) was performed for feature selection. Five-fold cross-validation was used to determine the tuning parameter for LASSO and LASSO feature selection was performed 100 times. Features that were selected 50% of the repetitions were used in the PLS-DA model. Model performance was determined by cross-validation.

### Safety and ethics

Experiments and procedures involving mice were approved under protocol LVD29E by the NIAID Animal Care and Use Committee according to standards set forth in the NIH guidelines, Animal Welfare Act, and US Federal Law. Euthanasia was carried out using carbon dioxide inhalation in accordance with the American Veterinary Medical Association Guidelines for Euthanasia of Animals (2013 Report of the AVMA Panel of Euthanasia). All procedures with infectious MPXV were performed in registered BSL-3 and ABSL-3 laboratories by trained and smallpox vaccinated investigators using protocols approved by the NIH Institutional Biosafety Committee.

## Acknowledgement

We would like to thank Rachel Leeson for her support in manuscript preparation. The technical staff of the NIAID Comprehensive Medical Branch provided excellent animal care. P.L.E, J.L.A and B.M. were supported by the Division of Intramural Research of NIAID. We would like to acknowledge USAMRIID’s Virology and Diagnostics Division personnel for contributing insights, reagents for assay development, and corroborating immune response data related to the studies reported herein.

## Author Contribution

Conceptualization: AWF, JWH, TC, AC, NJS, BM

Methodology & Investigation: AWF, CA, PLE, JLA, GYC, HN, TRF, JIM, GAA, CO, AN, MAD

Visualization: CA, GYC, AN, MAD

Project administration: HB, JJ

Supervision: GSJ, TC, AC

Writing – original draft: AWF, CA, GA, BM

## Data and materials availability

All data will be available as a supplemental table.

**Supplemental Figure 1. Sequence alignment and structural-divergence**..

a-d Sequence alignments of VACV-WR WT proteins versus MPXV-MA001 and MPXV Zaire 79 orthologous proteins for A27 (A), A33 (B), B5 (C) and L1 (D). Amino acid substitutions/deletions are shown in white background.

e-h Alpha-fold predicted 3D structures of MPXV-MA001 vaccine antigens used in this study are shown separately for A29 (E), A35 (F), B6 (G), and M1 (H). Mutations are shown in red spheres and numbering is relative to VACV-WR WT protein length (see alignments on the left).

**Supplemental Figure 2: Expression analysis of mRNA membrane-bound and secreted designs for monkeypox antigens**.

a Flow cytometry analysis of B6, A35 and M1 membrane-bound constructs indicating frequency of positive cells (left panel) and antigen surface-density represented by MFI*frequency (right panel) at indicated mRNA doses. In all cases an improvement in antigen surface density was observed compared to the viral reference sequences included for comparison.

b Immunoassay data for the selected A29 antigen sequences. The expected antigen molecular weight was observed at ∼14 kDa. We also observed the dimer at ∼28kDa in addition to an intermediate antigen-specific band at ∼21kDa.

**Supplemental Figure 3. Antibody isotype, subclass and FcR-binding against MPXV**.

a The dot plots show the total IgA, IgM, and IgG subclass response, as measured by a multiplex Luminex assay, against the antigen listed. The dotted line represents the median response in the PBS vaccinated group. Significance was determined by a Mann-Whitney U test and corrected for multiple hypothesis testing using Benjamini-Hochberg method. * p < 0.05, ** p <0.01, *** p<0.001, ****p<0.0001.

b The dot plots show the FcgR-binding response, as measured by a multiplex Luminex assay, against the antigen listed. The dotted line represents the median response in the PBS vaccinated group. Significance was determined by a Mann-Whitney U test and corrected for multiple hypothesis testing using Benjamini-Hochberg method. * p < 0.05, ** p <0.01, *** p<0.001, ****p<0.0001.

**Supplemental Figure 4. Univariate VACV-specific Luminex analysis**.

a The dot plots show the total IgA, IgM, and IgG subclass response, as measured by a multiplex Luminex assay, against the antigen listed. The dotted line represents the median response in the PBS vaccinated group. Significance was determined by a Mann-Whitney U test and corrected for multiple hypothesis testing using Benjamini-Hochberg method. * p < 0.05, ** p <0.01, *** p<0.001, ****p<0.0001.

b The dot plots show the FcgR-binding response, as measured by a multiplex Luminex assay, against the antigen listed. The dotted line represents the median response in the PBS vaccinated group. Significance was determined by a Mann-Whitney U test and corrected for multiple hypothesis testing using Benjamini-Hochberg method. * p < 0.05, ** p <0.01, *** p<0.001, ****p<0.0001.

