## Supplementary figures and images for "A monkeypox mRNA-lipid nanoparticle vaccine targeting virus binding, entry, and transmission drives protection against lethal orthopoxviral challenge"

### Supplemental Figure 1

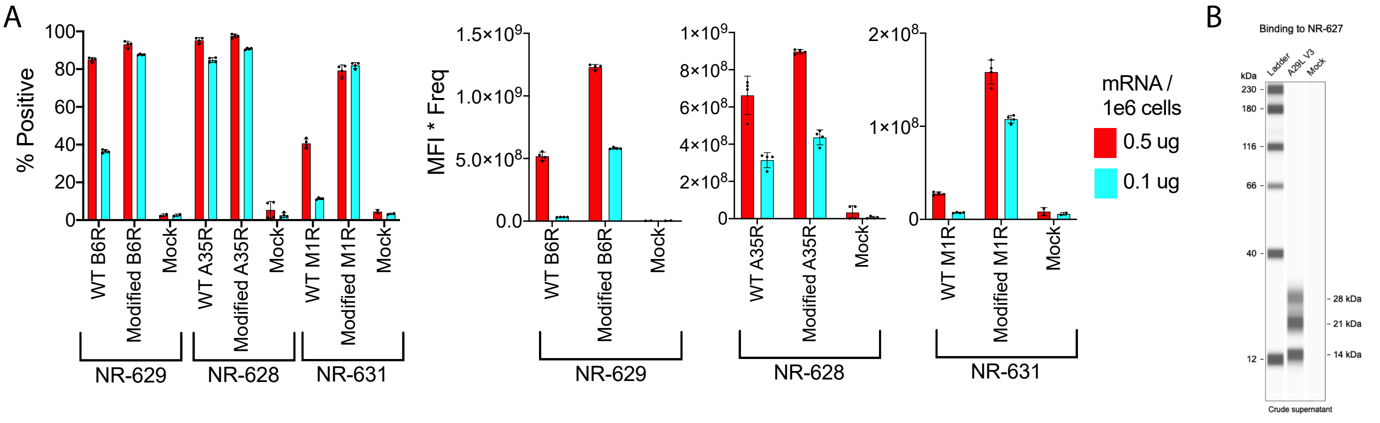

### Supplemental Figure 2

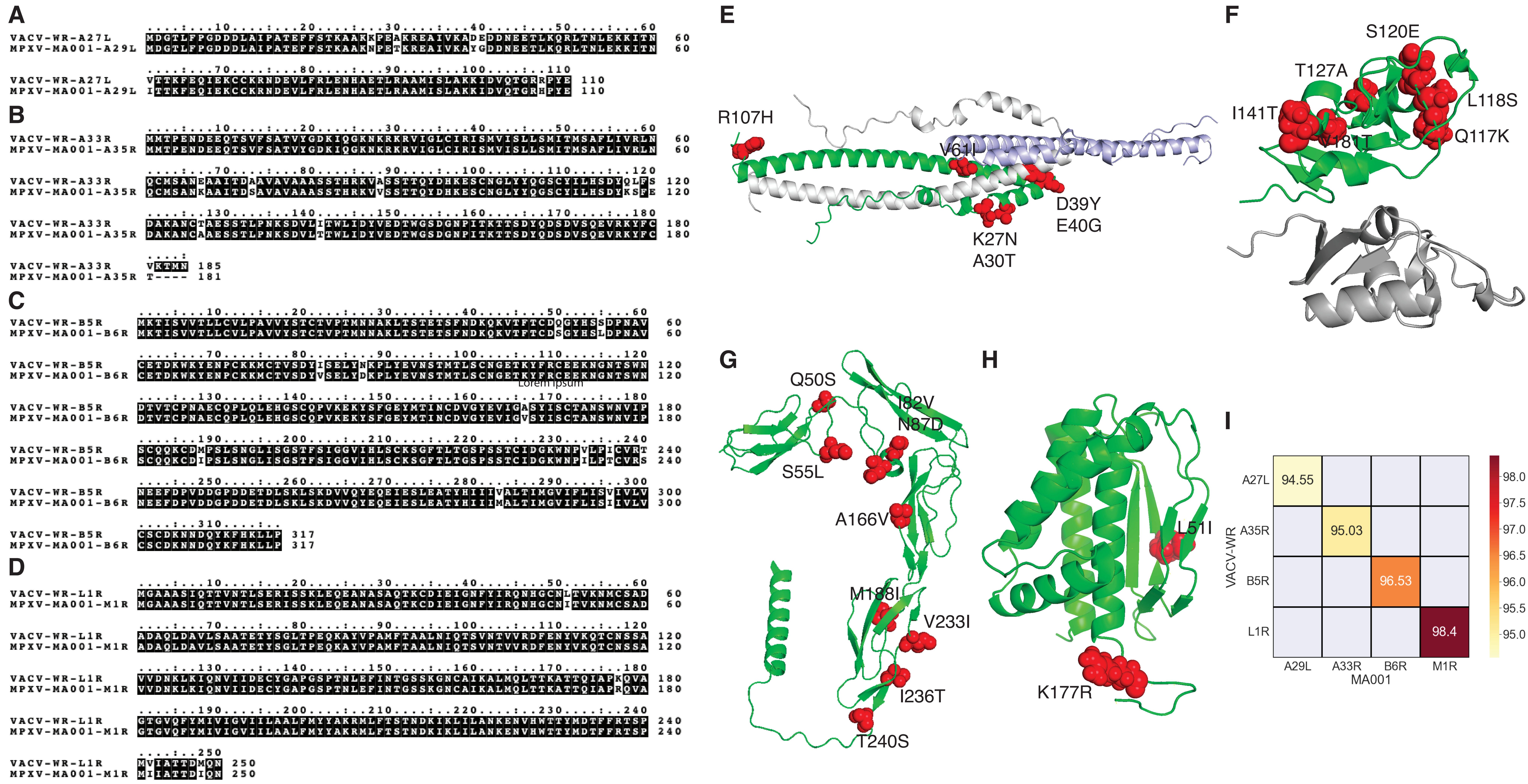

### Supplemental Figure 3

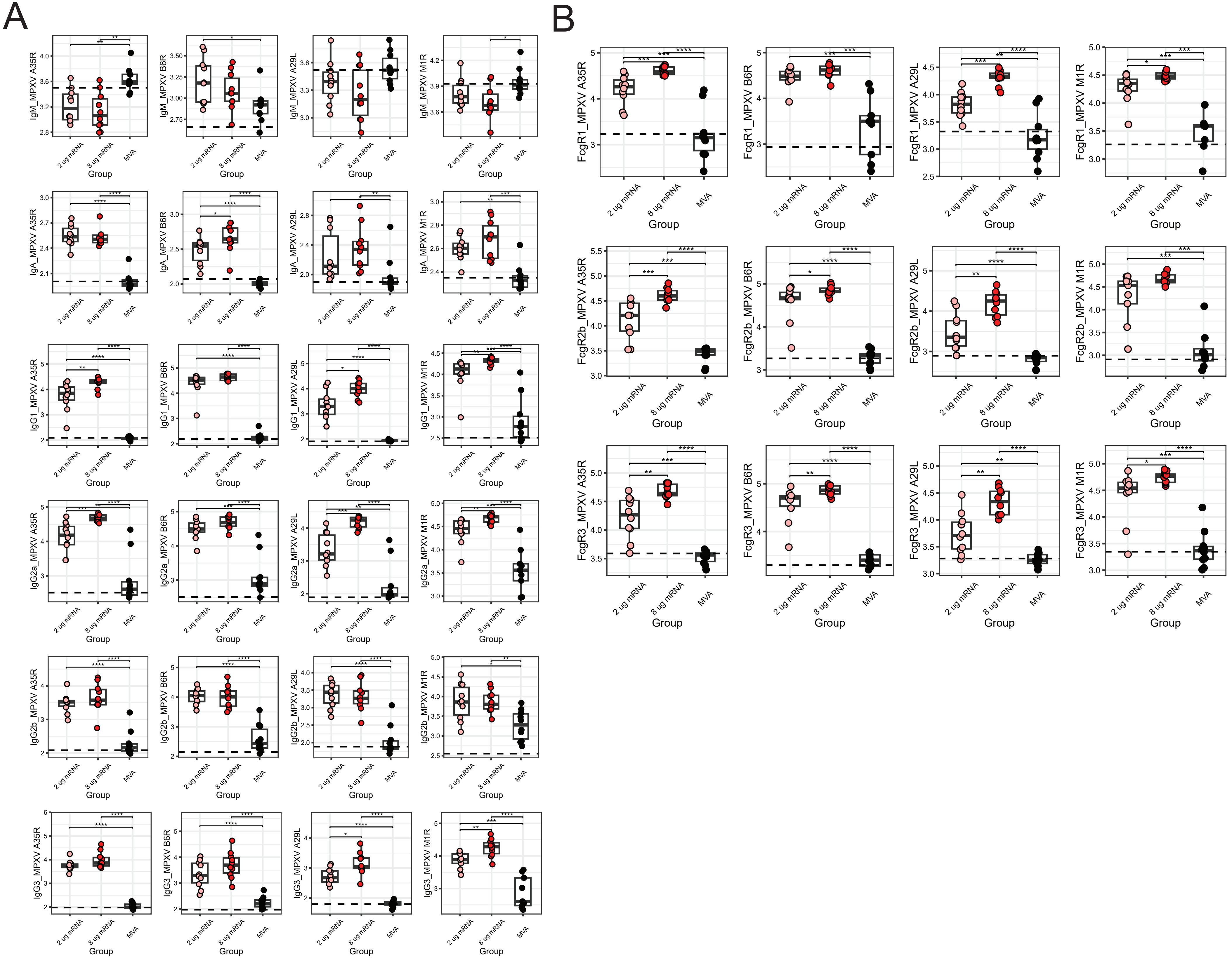

### Supplemental Figure 4

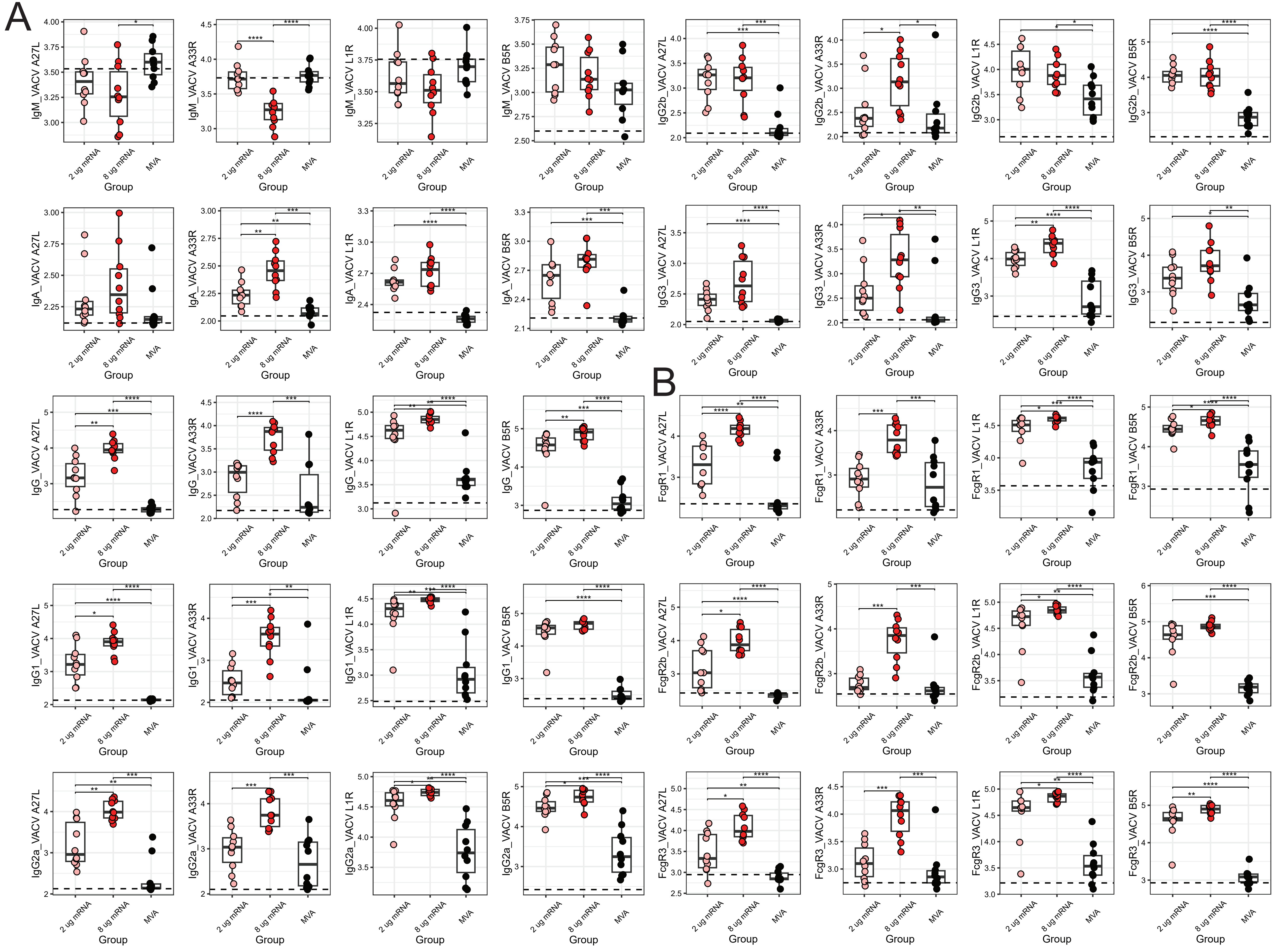
